# Prevalence of white matter pathways coming into a single diffusion MRI voxel orientation: the bottleneck issue in tractography

**DOI:** 10.1101/2021.06.22.449454

**Authors:** Kurt G Schilling, Chantal M.W. Tax, Francois Rheault, Bennett Landman, Adam Anderson, Maxime Descoteaux, Laurent Petit

## Abstract

Characterizing and understanding the limitations of diffusion MRI fiber tractography is a prerequisite for methodological advances and innovations which will allow these techniques to accurately map the connections of the human brain. The so-called “crossing fiber problem” has received tremendous attention and has continuously triggered the community to develop novel approaches for disentangling distinctly oriented fiber populations. Perhaps an even greater challenge occurs when multiple white matter bundles converge within a single voxel, or throughout a single brain region, and share the same parallel orientation, before diverging and continuing towards their final cortical or sub-cortical terminations. These so-called “bottleneck” regions contribute to the ill-posed nature of the tractography process, and lead to both false positive and false negative estimated connections. Yet, as opposed to the extent of crossing fibers, a thorough characterization of bottleneck regions has not been performed. The aim of this study is to quantify the prevalence of bottleneck regions. To do this, we use diffusion tractography to segment *known* white matter bundles of the brain, and assign each bundle to voxels they pass through and to specific orientations within those voxels (i.e. fixels). We demonstrate that bottlenecks occur in greater than 50-70% of fixels in the white matter of the human brain. We find that all projection, association, and commissural fibers contribute to, and are affected by, this phenomenon, and show that even regions traditionally considered “single fiber voxels” often contain multiple fiber populations. Together, this study shows that a majority of white matter presents bottlenecks for tractography which may lead to incorrect or erroneous estimates of brain connectivity or quantitative tractography (i.e., tractometry), and underscores the need for a paradigm shift in the process of tractography and bundle segmentation for studying the fiber pathways of the human brain.

## Introduction

> *Two paths diverged from a single orientation*,
>
> *And streamlines could not travel both*
>
> *Else it be a false positive, long it stood*
>
> *And looked down one as far it could*
>
> *To which cortex should it approach?*

Diffusion MRI fiber tractography is currently the only tool to map the long-range structural brain connectivity *in vivo*. However, there are a number of limitations and ambiguities that affect the ability of tractography to accurately map the connections of the brain [1]. At the *voxel level*, significant attention has been given to the “crossing fiber problem” [2–4]. This problem typically refers to the situation when two or more *differently oriented* fiber bundles are located in the same imaging voxel, which causes a partial volume effect that can lead to ambiguous or incorrect estimates of fiber orientation and subsequent failure of tractography [5]. Crossing fibers have been shown to occur in a majority of the voxels in the brain [6], and for the last decade the crossing fiber problem has been cited as the major limitation that diffusion tractography faces, with a vast number of algorithms [7] and papers referring to this problem [1–4, 8–24]. The identification and characterization of this problem led to a fundamental paradigm shift in diffusion processing, moving the field beyond classical diffusion tensor imaging, and has led to the development of a number of algorithms capable of resolving crossing fibers [2–4, 13, 25, 26].

A more recently described limitation of fiber tractography is the “bottleneck problem” [27]. In contrast to the crossing fiber problem, bottlenecks occur at a *global level* when multiple fiber populations converge towards a narrow region, temporarily aligning and sharing the *same orientation* and trajectory, before re-emerging from the bottleneck region [1, 14]. Current tractography algorithms cannot adequately choose the correct pathway upon reemerging, which leads to generation of a potentially large number of false positive pathways [28], and limits the ability to use tractography as a tool to explore potential connections and fiber pathways of the brain. While significant efforts have gone into solving the crossing fiber problem, the bottleneck problem has received far less attention. A thorough characterization and investigation of bottleneck locations and prevalence may highlight the extent of this problem, and much like the crossing fiber problem, cause a paradigm shift in tractography in order to solve this issue.

In this work, we utilize well-known and well-characterized white matter fiber bundles extracted using automated tools, to quantify how often they overlap within the same imaging voxels, and also how often the overlap occurs within the same voxel while also sharing the same dominant orientation. These locations represent known bottleneck regions for tractography, and indicate areas in the brain where a number of white matter pathways converge, and where tractography may lead to incorrect or erroneous estimates of brain connectivity.

### Nomenclature

Here, we aim to clarify nomenclature that will be used in this study to describe our methodology and results. First, a **bundle**, or fiber bundle, is a group of streamlines that is created from a diffusion MRI dataset and is intended to represent a specific white matter pathway of the brain. Bundles contain streamlines with start and end points generally belonging to the same brain territories, respectively. In this study, we create bundles using two common white matter atlases and tractography dissection techniques that are informed by prior anatomical knowledge and contain pathways for which there is extensive evidence of their existence. Thus, in this study we are analyzing only *anatomically plausible* bundles.

Next, a **voxel** represents orientational information in threedimensional space. In MRI, the size of voxels is on the order of millimeters, whereas axons of the brain have diameters on the scale of micrometers, and a single voxel can contain hundreds of thousands of axons. The directionality of axons within a voxel can be summarized by the **fiber orientation distribution (FOD)**. The FOD is a continuous function over a sphere and can be visualized as a 3D directional histogram where peaks, or local maxima, are assumed to point parallel to the direction of axons. The FOD can be segmented into discrete elements based on peaks, or lobes, that are considered to be representative of a specific orientation of a set of axons within each voxel. These fiber elements are referred to as **fixels** [29], and are parameterized by the mean orientation of fibers within the lobe of the FOD. Thus, there can be multiple fixels in a single voxel, with the advantage that we can now assign a specific property or index to each fixel, or orientation, within a voxel.

For this study, it is necessary to clarify or define six classifications, written out in Table 1 and displayed as a cartoon in **Figure 1**. If a FOD has only a single peak, or local maxima, we classify it as a **single-orientation (fiber) voxel**, and if it has greater than one peak we call this a **multiorientation voxel** [6]. In contrast to simply counting peaks, we also count the number of bundles passing through the same voxel, then characterizing as a **single-bundle voxel** and **multi-bundle voxel**. Similarly, we count the number of bundles associated with each fixel and, characterize the fixel as a **single-bundle fixel**, or **multi-bundle fixel**. As described above, a fixel is usually used to describe a single fiber bundle element. However, we hypothesize that a single fixel may be associated with several bundles, thus creating bottlenecks for tractography. **A bottleneck ‘region’** then is a cluster of coherent multi-bundle fixels.

**Figure 1.**
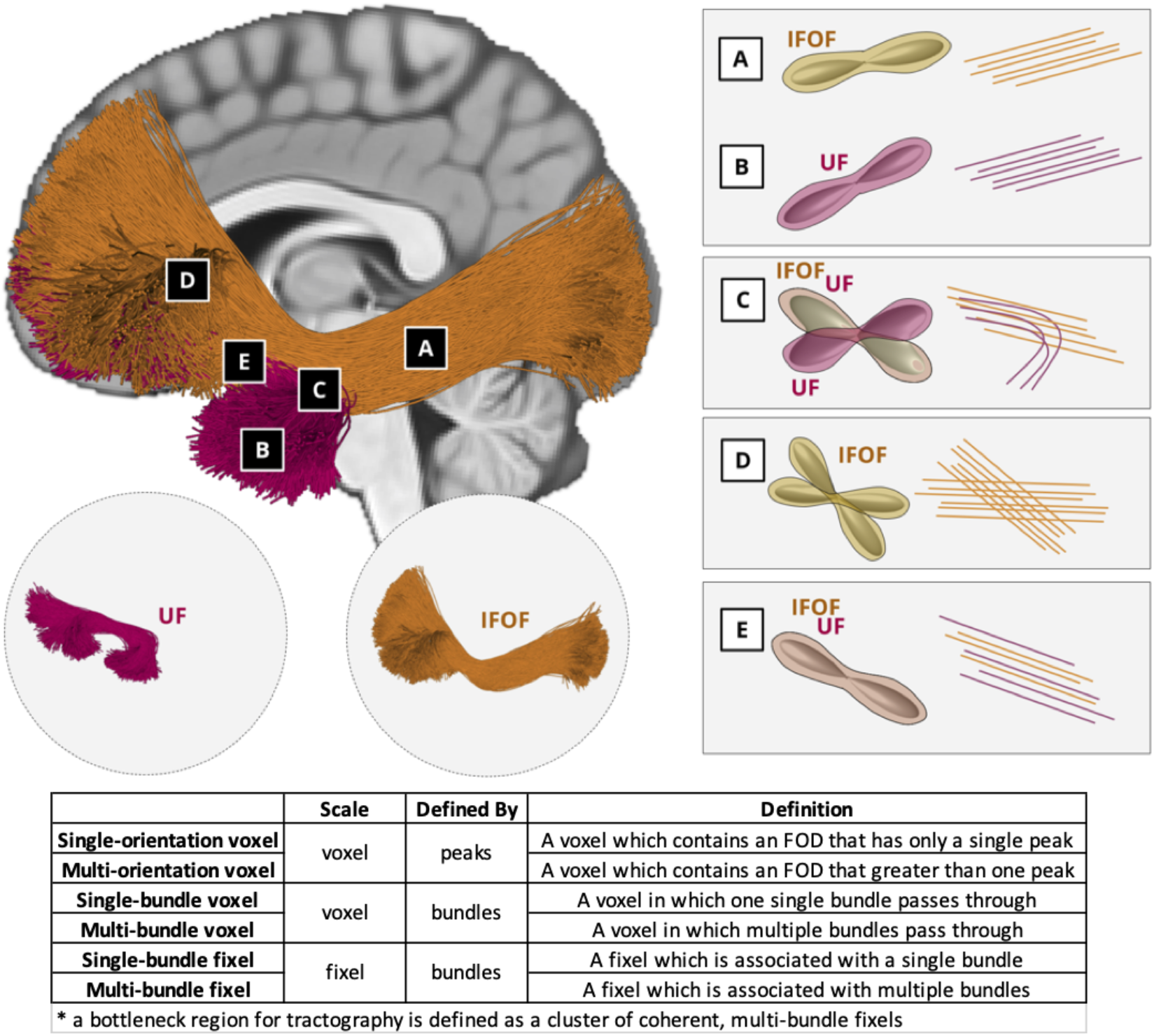
Nomenclature. Two bundles, the UF and IFOF, are used to highlight classifications of voxels (A-D), and fixels within the voxels. Voxels in A and B are examples of single-orientation voxel and single-bundle voxels and also single-bundle fixel. Because the UF and IFOF diverge in Voxel C, this is an example of a multi-orientation voxel and multi-bundle voxel, with one fixel classified as a single-bundle fixel and the other a multibundle fixel. Voxel D highlights the fanning of the IFOF, which results in a multi-orientation voxel and single-bundle voxel, and both fixels are single-bundle fixels. Finally, both the IFOF and UF pass through voxel E following the same orientation, thus voxel E is a single-orientation voxel but multi-bundle voxel, and also a multi-bundle fixel. This fixel, and thus also this voxel, represents a bottleneck for tractography.

## Methods

### Data

We utilized data from 25 healthy subjects in the Human Connectome Project (HCP) S1200 release [30]. The HCP protocol (custom 3T Siemens Skyra) included T1-weighted images acquired using a 3D MPRAGE sequence (TE = 2.1 ms, TR = 2400 ms, flip angle = 8 deg, FOV = 224×224mm, acquisition, voxel size = 0.7mm isotropic). Diffusion images were acquired using a single-shot EPI sequence, and consisted of three b-values (b = 1000, 2000, and 3000 s/mm^2^), with 90 directions (and 6 b=0 s/mm^2^) per shell (TE = 89.5 ms, TR = 5520 ms, slice thickness = 1.25 mm, flip angle = 78 degrees, FOV = 210*180, voxel size = 1.25mm isotropic). Data pre-processing included correction for susceptibility distortions, subject motion, and eddy current correction.

### Processing

All processing was performed with the MRTrix3 software package [31]. Multi-shell, multi-tissue constrained spherical deconvolution (dwi2fod) was used to estimate the white matter FOD for each subject using a group averaged white matter response function computed using the “dhollander” algorithm [32]. Next, we generated a study-specific unbiased FOD template (population_template) for population-based analysis.

From both the population FOD template and subject-specific FODs, we extracted peaks (sh2peaks) and fixels (fod2fixel), with a threshold of 0.1 times the maximum peak amplitude to remove spurious peaks. We then counted the number of peaks (i.e., lobes of the FOD) and defined those voxels which have only a single peak to be a **single-orientation voxel**, and those with greater than one peak to be **multi-orientation voxels**.

### Bundle segmentation

We utilized two common, automated, pipelines for white matter bundle extraction, Recobundles [33] and TractSeg [34, 35]. These are both informed by prior anatomical knowledge in order to generate bundles representative of well-characterized, and well-validated, white matter pathways of the brain. All analysis is performed separately for both techniques to show that results generalize across slight deviations in the number and definitions of white matter pathways, and the techniques used to extract them.

Recobundles is based on an atlas of 78 bundles [36], although 12 bundles are cranial nerves outside of the cerebrum and brainstem. Whole brain tractography was performed using anatomically constrained probabilistic tractography with the iFOD2 propagation algorithm (tckgen), to generate 25 million streamlines, which were filtered based on the diffusion signal using the SIFT algorithm [37] (tcksift) to 2 million streamlines. Bundle recognition was performed following streamline linear registration to the HCP842 [36] bundle template (dipy_slr) and bundle recognition using default parameters of the RecoBundles algorithm [33].

TractSeg [35] is a tool based on convolutional neural networks that is trained to create tract orientation maps and segmentations of end regions, which can be used to perform probabilistic bundle-specific tractography [34]. We implemented the processing pipeline provided at (https://github.com/MIC-DKFZ/TractSeg) with the MRTrix-derived FODs as input, in order to generate 72 bundles per subject.

### Assigning bundles to voxels and fixels

**Figure 2** visualizes the procedure used to assign bundles to voxels and bundles to fixels. For each bundle (**Figure 2, A**; N=78 RecoBundles; N=72 TractSeg), a tract density map was created (tckmap) by counting the number of streamlines coursing through each voxel (**Figure 2, B**). Density maps were thresholded at 5% of the maximum streamline density per bundle in order to create binary segmentations indicating the voxel-wise profile of each bundle. Next, a fixel density map (**Figure 2, C**) was created (tck2fixel) by counting the number of streamlines that are most closely aligned to each fixel. Again, fixel-density maps were thresholded at 5% of the maximum density in order to create binary segmentations indicating the fixel-wise profile of each bundle. Now, the number of known bundles per voxel, as well as number of known bundles per fixel, can be counted. A single fixel that is associated with multiple bundles represent a potential bottleneck. Voxel-based and fixel-based analysis is performed both at the level of the individual in subject-space, and also at the population-level in template-space.

**Figure 2.**
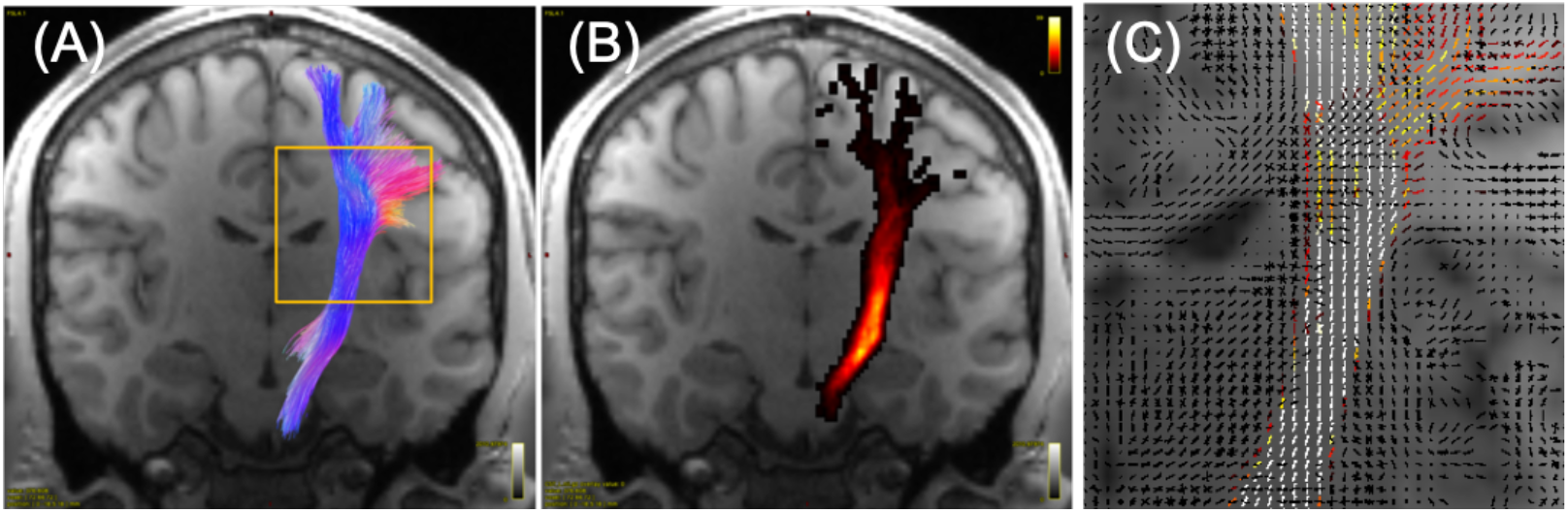
Assigning bundles to voxels and fixels. Each segmented white matter bundle (A) was assigned to each voxel (B) based on thresholding of the density map, as well as assigned to each fixel (C) within a voxel based on matching orientations of streamlines. This allowed us to query both the number of known bundles per voxel, and number of known bundles per fixel. White matter bundles were derived from TractSeg (N=72 bundles) and Recobundles (N=78 bundles).

## Results

### Bottleneck Prevalence

Investigating fixels throughout the white matter, we first ask *what is the prevalence of multi-bundle fixels*? **Figure 3** shows maps of the number of bundles assigned to each fixel, visualized in coronal, sagittal, and axial views, for both TractSeg bundles (top) and Recobundles (bottom). Most noticeably, most fixels in the white matter, in all orientations, are associated with multiple bundles. In fact, many regions have groups of oriented fixels associated with 7+ unique bundles. **Figure 4** quantifies the number of bundles assigned to each individual fixel. The results confirm the qualitative observation that most fixels contain multiple bundles converging in a given orientation, with greater than 50% of fixels in Recobundles and greater than 70% of fixels in TractSeg containing greater than a single fiber population. In general, TractSeg bundles show a higher prevalence of multi-bundle fixels (i.e., bottlenecks) than Recobundles. In summary, a majority of fixels in the brain that contain known fiber bundles act as bottleneck regions for tractography.

**Figure 3.**
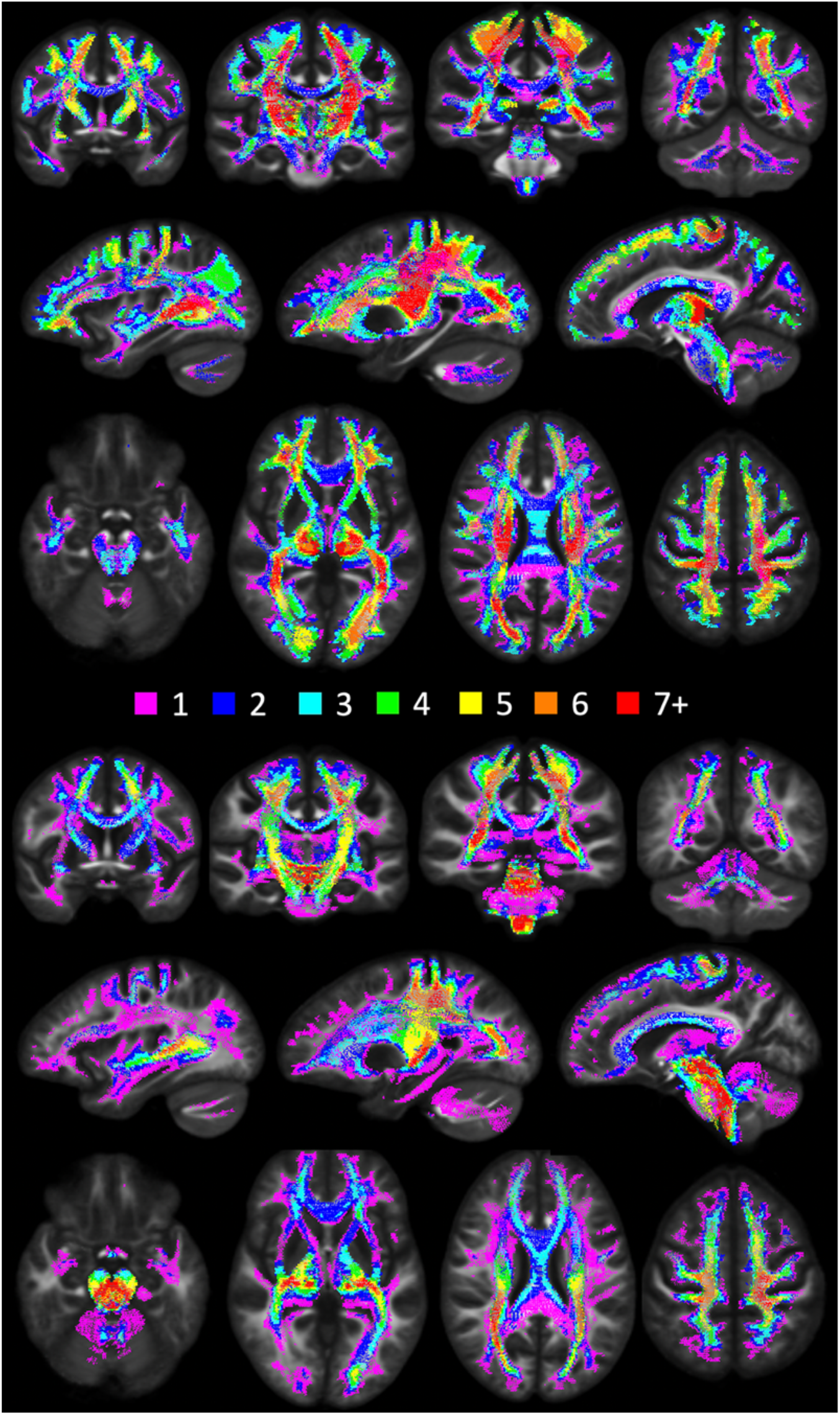
There is a high prevalence of bundles assigned to each fixel in the brain. Results are shown as vectors, colored by the number of associated bundles, and averaged across the population. TractSeg results are shown on top, Recobundles on bottom. Fixels with more than one bundle traversing through them represent bottleneck regions for tractography.

**Figure 4.**
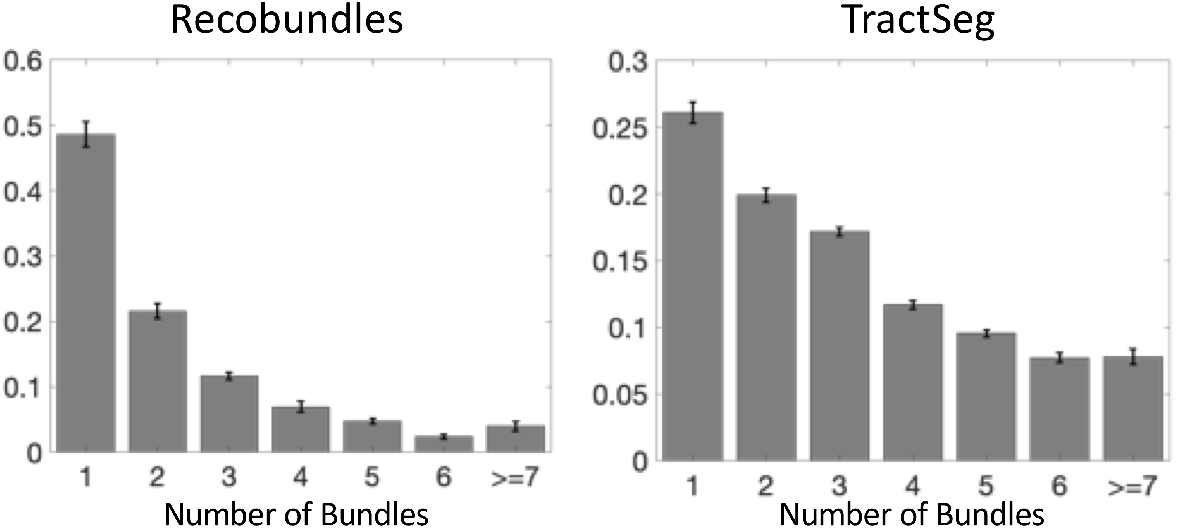
Number of bundles assigned to fixels in the brain. Most fixels had greater than one bundle traversing through their designated orientation. Note that fixels which were assigned to 0 bundles are not shown. Y-axis is shown as a fraction of fixels. Error bars represent variation across the studied population.

### Bottleneck Locations

Next, we ask *where do the most important bottleneck regions occur?* These are regions in which groups of fixels with similar orientation exhibit a convergence of the largest number of pathways. Highlighted bottleneck regions, e.g., 7 or more bundles, are visualized in **Figures 5–8**. In all Figures, the fixels are color-coded by the number of fibers traversing through each orientation, and exemplar bundles converging in each region are displayed.

**Figure 5.**
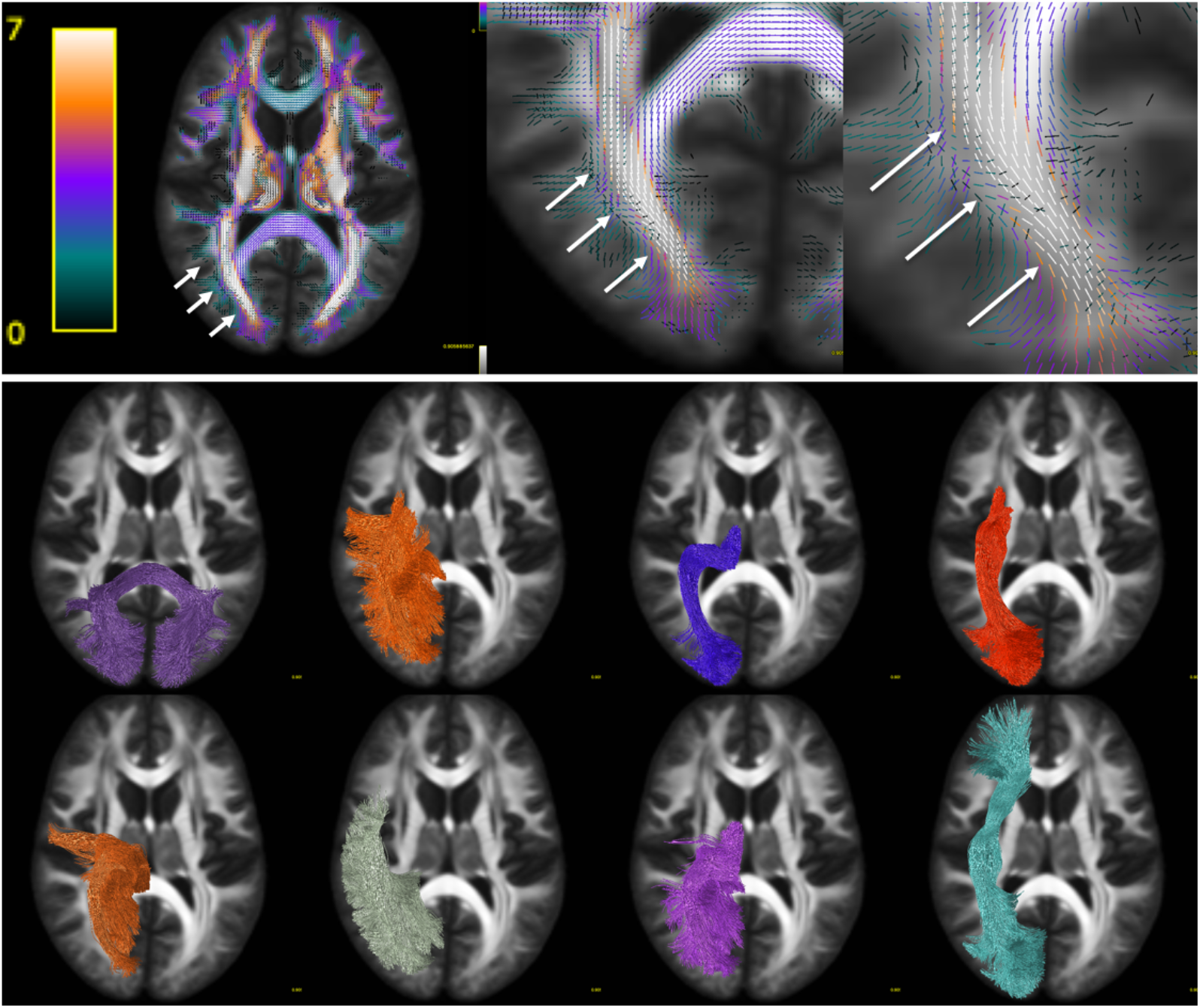
Bottleneck region in the anterior-posterior oriented white matter of the occipital lobe (arrows) contains a large number of white matter bundles with unique starting and ending connections. Colormap ranges from 0 to 7+ bundles. Pathways (derived from TractSeg) from left to right, top to bottom: CC7, ST_par, OR, ST_OCC, POPT, MdLF, T_PAR, IFO (for full names see acronyms at end of document).

**Figure 6.**
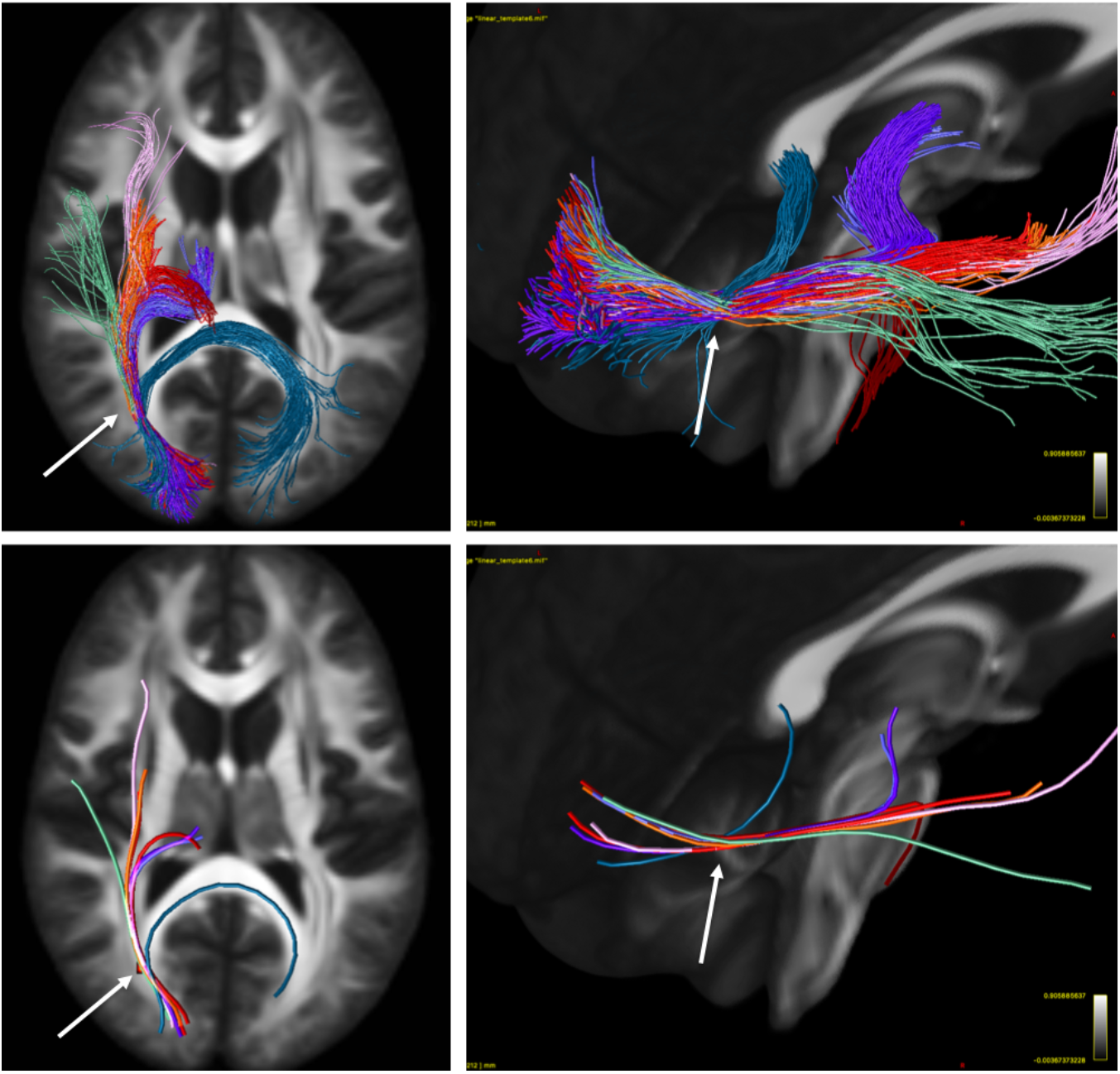
Illustration of the bottleneck in the anterior-posterior oriented white matter of the occipital lobe. Fiber bundles from Figure 5 (same color scheme) were filtered to select only streamlines (top) traversing a small 2×2×2 voxel region of interest (arrow). A single representative streamline from each sub-bundle is also shown (bottom). This example emphasizes that streamlines belonging to many fiber bundles may traverse through the same small region, in the same orientation.

**Figure 7.**
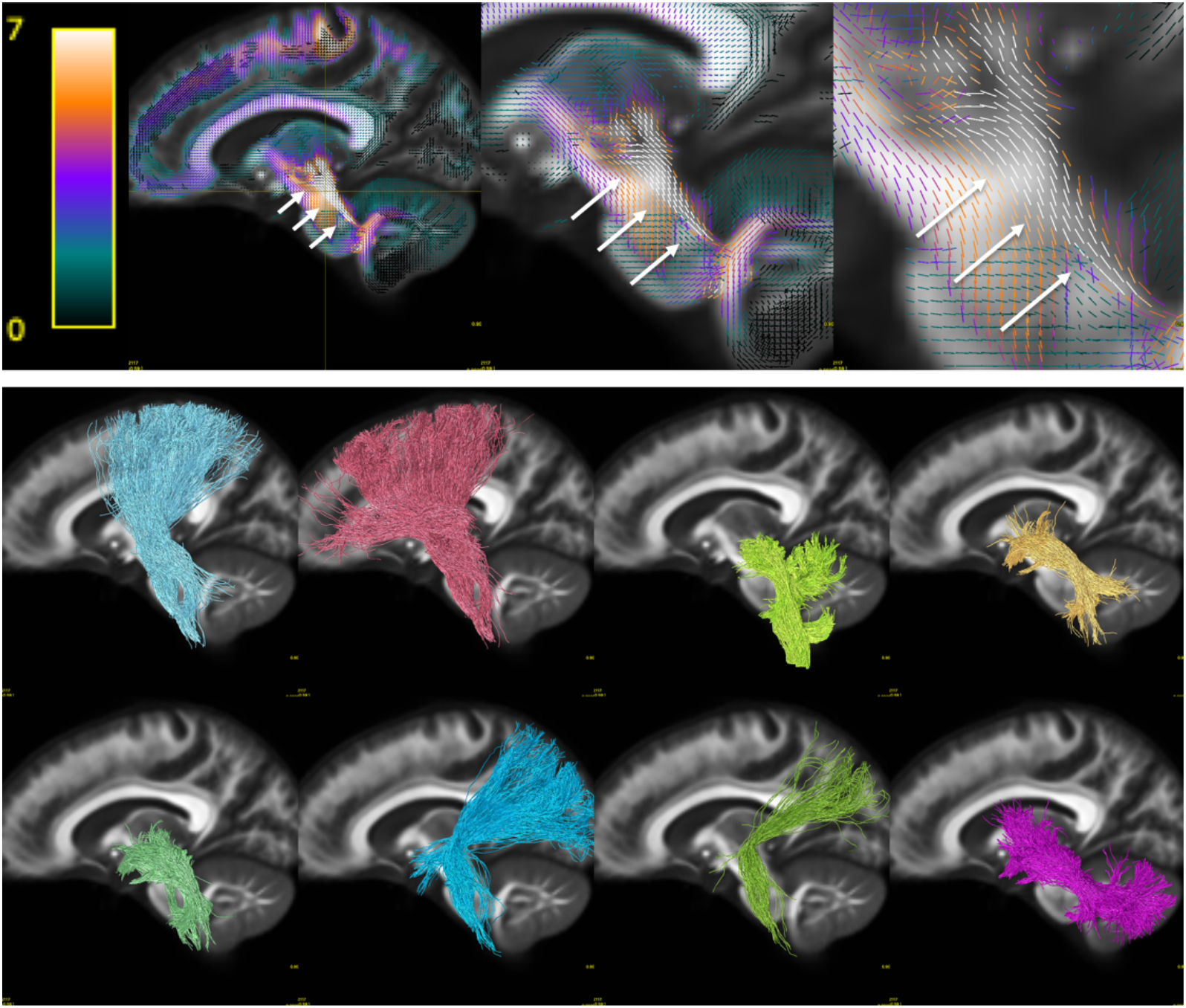
Bottleneck region in the superior-inferior oriented white matter of the brain-stem (arrow) contains a large number of white matter bundles with unique starting and ending connections. Hot-cold colormap ranges from 0 to 7 bundles. Pathways (derived from Recobundles) from top to bottom, left to right: CST, FPT, LL, MLL, CTT, OPT, TPT, SCP (for full names see acronyms at end of document).

**Figure 8.**
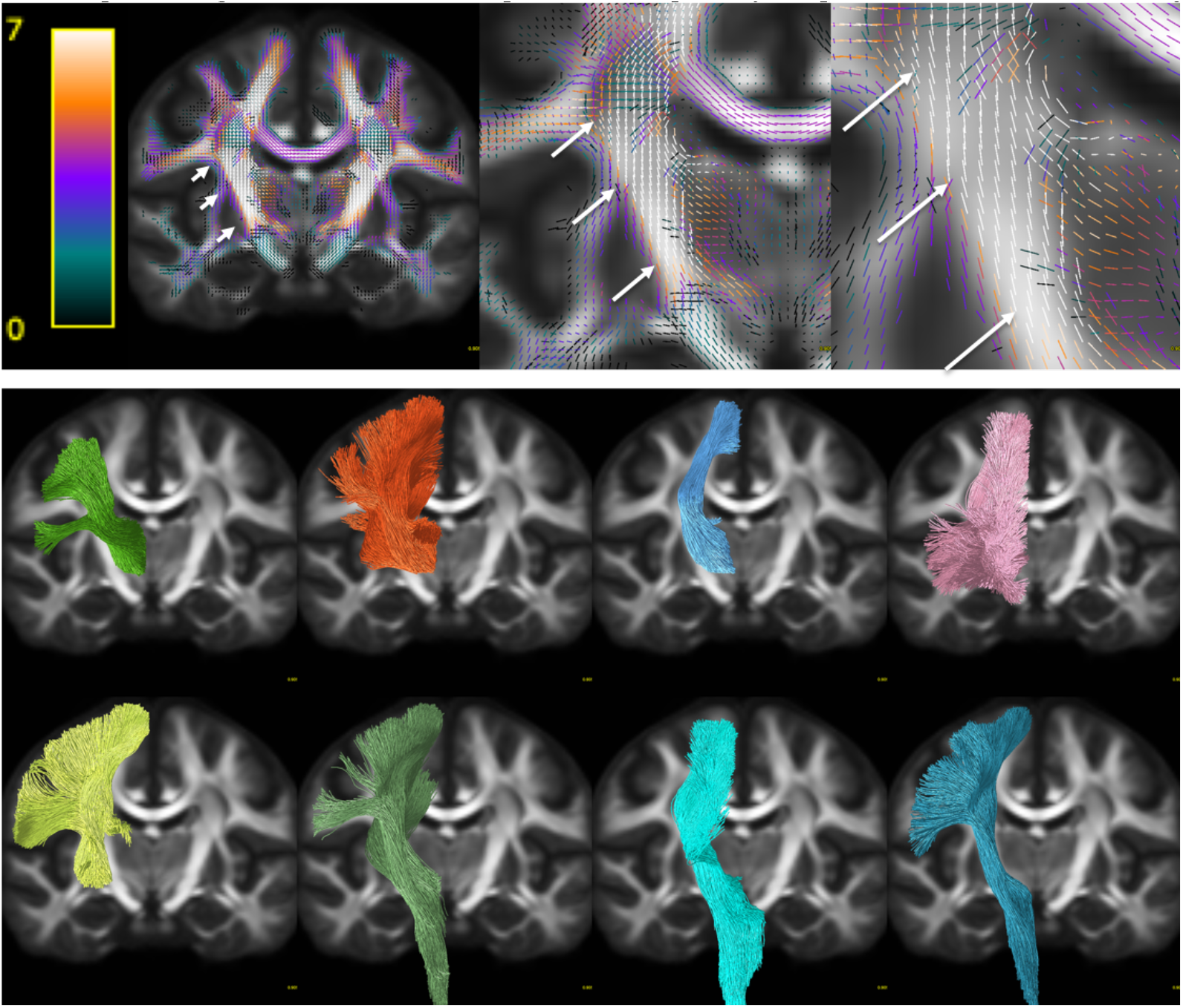
Bottleneck region in the superior-inferior oriented white matter of the internal capsule (arrow) contains a large number of white matter bundles with unique starting and ending connections. Hot-cold colormap ranges from 0 to 7 bundles. Pathways (derived from TractSeg) from left to right, top to bottom: T_PREM, T_PAR, STR, ST_PREF, ST_POST, POPT, FPT, CST (for full names see acronyms at end of document).

The first displayed region with the highest number of bottlenecks (**Figure 5**) is located in the deep white matter of the occipital lobe oriented in the anterior-posterior direction, covering a wide dorsal-ventral expanse and including the stratum sagittale. A large number of individual WM pathways, each with unique starting and/or ending connections, converge through this region with the same orientation, including the posterior part of the inferior fronto-occipital fasciculus (IFOF), inferior longitudinal fasciculus (ILF), middle longitudinal fasciculus (MdLF), parieto-thalamic and occipito-thalamic (optic radiations, OR) connections, parieto-striatal and parieto-occipital pontine tract (POPT), and several subdivisions or segmentations of the striato-cortical connections, and splenium of the corpus callosum. Thus, while most pathways terminate throughout the occipital lobe, these fibers can project to sub-cortical nuclei, temporal or frontal lobes, or to the contralateral hemisphere as commissural fibers.

**Figure 6** further highlights the bottleneck problem in this region, and illustrates how different tracts may share a similar location AND orientation, yet have unique start and end points. We filtered the previously described bundles using a single region of interest (a 2×2×2 voxel cube i.e., a 2.5mm isotropic region), and show just the streamlines from each bundle that traverse this area (top row). While the full extent of each pathway does not traverse this region, a large and coherent subset of each bundle does, all oriented in the anterior-posterior direction. To simplify the illustration, a representative streamline is shown for each filtered bundle (bottom row), exemplifying the bottleneck problem: there is a large combinatorial number of possible pathways that traverse through this voxel following this single well-defined orientation.

The second main bottleneck region is the convergence of superior-inferior oriented fibers converging and traversing throughout the brainstem, from the mid-brain to the medulla (**Figure 7**). This includes a number of ascending and descending fibers projection pathways, corticopontine fibers arising from the cortex, and cerebellar tracts. Again, fibers traversing through this region all share the same dominant orientation, yet end throughout the extent of the cortex, subcortex, spinal cord, and cerebellum.

The third region with highest number of bottlenecks occurs in the superior-inferior oriented fibers of the posterior limb of the internal capsule (**Figure 8**). This region contains pathways such as the corticospinal tract, frontal and parietal pontine fasciculi, striato-postcentral and striato-precentral bundles, superior thalamic radiations toward the parietal, precentral, and postcentral cortices. While most of these fibers project from the mid-brain and nuclei, projections cover the expanse of the parietal and frontal cortices, with many projecting onward dorsally towards the superior frontal gyrus.

### Single-bundle and multi-bundle voxels

Rather than assessing fixels, we can also ask *what is the prevalence of single-bundle and multi-bundle voxels*, and *where do known bundles overlap?* Thus, this overlap can contain both bundles oriented in the same direction, and different directions within a voxel. **Figure 9** quantifies the number of bundles per voxel for both Recobundles and TractSeg bundles, and visualizes bundle overlap in the template space. It is clear that a majority of voxels in the white matter contain multiple, overlapping bundles. In fact, a number of voxels in the brain again contain as many as 7 or more unique overlapping bundles, with overlap of these bundles frequently occurring in regions that parallel the bottleneck regions - the centrum semiovale and posterior corona radiata, and also the posterior limb and retrolenticular limb of the internal capsule. Thus, most white matter voxels (>65-75%) contain overlap of multiple bundles.

**Figure 9.**
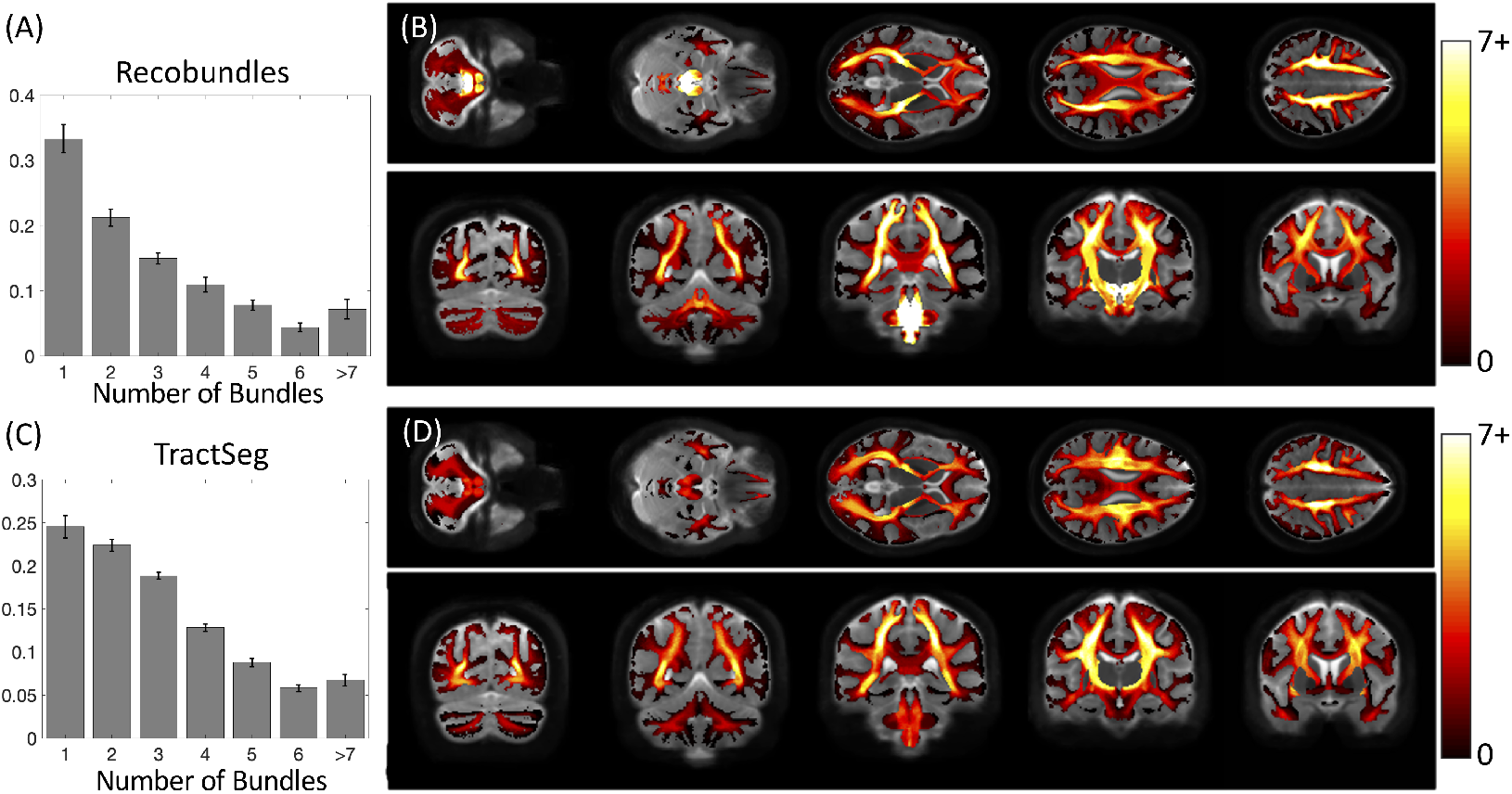
Many voxels in the white matter contain multiple known bundles. Prevalence of voxels with 1 to 7+ bundles is quantified for Recobundles (A) and visualized in template space (B), and also quantified for TractSeg bundles (C) and visualized overlaid in template space (D). Note that many voxels contain 0 bundles (i.e, are not associated with known bundles in our atlas) and are thus not quantified.

### Single-orientation and multi-orientation voxels

Finally, we ask *what is the prevalence of single-orientation and multi-orientation* voxels (i.e., the prevalence of the “crossing fiber problem”), and *where do single- and multiorientation voxels occur?* **Figure 10** quantifies and visualizes the prevalence of multi-orientation voxels in both an individual subject and across the population. In agreement with previous literature, our results show that a majority of voxels in the white matter (>60%) contain multiorientation voxels. These voxels are prevalent throughout the entire white matter, with more complex (for example >2 peaks) crossings in the centrum semiovale and cerebellum. Voxels with only a single peak are prevalent in the corpus callosum and internal capsule, as well as near the crowns of various gyri.

**Figure 10.**
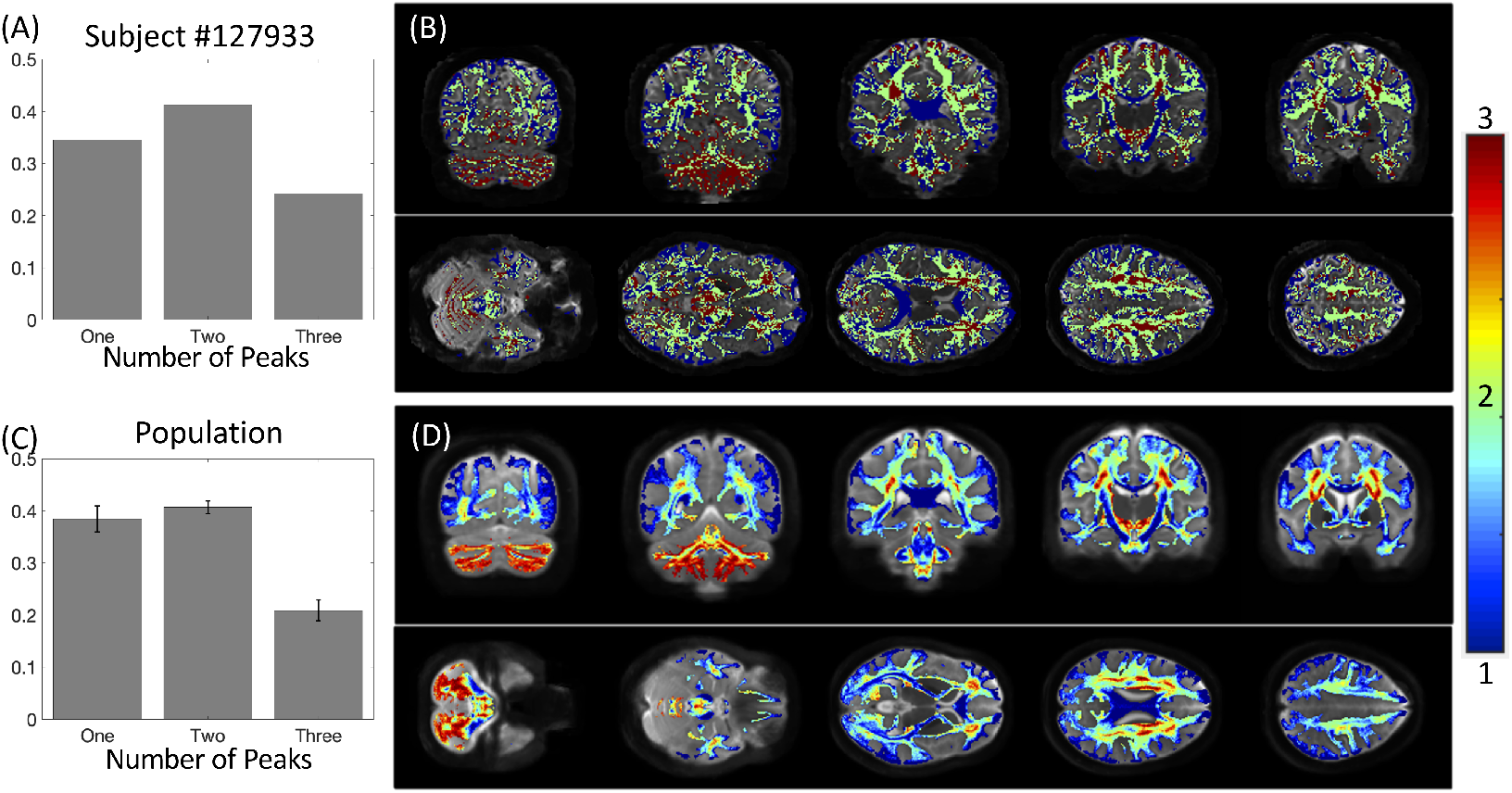
There is a high prevalence of multi-orientation voxels throughout the brain. Prevalence of voxels with one peak, two peaks, and three peaks is quantified for a single subject (A) and visualized overlaid on an anatomical image (B) and also averaged across the population (C) and visualized overlaid across the population template (D). Note that the number of peaks is discrete on a single subject but continuous when averaged across all subjects.

One particularly interesting finding is that many of the identified bottlenecks above are often associated with single-orientation voxels - the internal capsule, mid-brain, and less frequently in the deep white matter of the occipital lobe.

## Discussion

In this study, we investigated the prevalence and locations of bottleneck regions in the brain, which present obstacles in our ability to build anatomically correct maps of the human brain using diffusion tractography.

We find that not only do a majority of voxels contain multiple bundles, but also that a majority of individual FOD peaks within a voxel (i.e., fixels) are associated with multiple bundles. As much as 50-70% of fixels in the brain contain multiple fiber bundles traversing through them. This is based on the use of “well-known” bundles, representing only a lower-bound that can only go up as we understand and map all existing bundles in the brain. The convergence of bundles into a nearly parallel funnel, and subsequent convergence, may lead to a combinatorial number of possible pathways that tractography algorithms may choose to take, leading to the generation of possibly anatomically non-existent pathways. While this may not present a problem for bundle-specific tractography with the use of manually placed priors [38], bundle templates [39, 40], or machine learning [35, 41, 42], mapping the entirety of the human connectome ultimately strives to map *all* of the unique bundles of the brain. The identified bottleneck regions clearly present obstacles to this mapping when the true connections are not known *a priori*. From a whole-brain connectome, streamlines which pass through these fixels without explicit knowledge of their existence should be suspected to be false positive connections generated by the process. Over and above problems created in connectomics studies, this also highlights the problems of proposing the existence of a new or unique pathway from diffusion MRI alone, even if the pathway is reproducible across scans and subjects. We posit that orthogonal information in the form of blunt dissection, tracers in animals, and alternative contrasts is necessary for inferences which demand highly specific tractography.

This bottleneck problem has further implications on quantitative tractography, or *tractometry*, where the along-tract profile of measures along multiple tracts allows a comprehensive characterization of white matter. Most methods map voxel-wise values along the tract (such as FA) [43], which we know are affected by crossing fibers, while recent studies map fixel-wise values (such as apparent fiber density), along the tract [44, 45]. Yet, we show here that these measures are still not yet truly specific to one bundle. Finally, studies that use global methods of filtering or quantification [37, 46] can map an average streamline-specific estimate of diffusion or relaxometry along the tract [47], however, these will still be affected by partial volume at different points along the streamline profile.

We additionally confirm findings from previous studies which indicate that most voxels have multiple peaks in their FOD. Previous studies have estimated anywhere from 60-90% of the brain contains “crossing fibers” [19, 21], estimates which vary with signal to noise ratio, diffusion sensitivity, and image resolution [6, 9]. Our estimated fraction is at the low-end of literature values, but we utilize multi-shell deconvolution as opposed to previous work, which may result in more regularized reconstruction and less spurious peaks. Regardless, this suggests that most *voxels* have more than one bundle traversing its location. Over and above this, we find that most *peaks within a voxel* have more than one bundle traversing in its direction. Thus, even solving the crossing fiber problem does not solve the bottleneck problem that may cause a large number of ambiguous, false positive pathways. Here, we propose that characterizing and describing the prevalence of this problem should lead to the development of methods which may alleviate or overcome these obstacles, much like that done for the crossing fiber problem. It is likely that advances in anatomical knowledge in combination with innovation in streamline generation and innovation in streamline filtering, will be needed to mitigate this problem.

### Bottleneck locations

We have described the highest bottleneck regions in this study. Specifically, we highlighted the deep white matter of the occipital lobe, the brainstem, and the internal capsule. These regions included a number of association, projection, and commissural fibers, all with unique trajectories and fundamentally different structural connections. While these were the most visually apparent ‘hot spots’, it is clear that a majority of fixels in the brain are associated with multiple white matter fiber pathways. From a tractography perspective, these regions may cause ambiguous connectivity estimates, yet, anatomically, these locations may have significant functional relevance, representing the intersection or merging of the many anatomo-functional highways of the brain.

Importantly, these bottlenecks are almost certainly an underestimation of the true prevalence and extent of this problem. We choose segmentation techniques which reconstruct only known anatomical pathways of the brain (72 and 78 bundles, respectively) for which there is broad agreement on their existence. Several other segmentation techniques exist which suggest the existence of a much greater number of unique pathways in the brain, however we have chosen to perform a conservative estimation, in addition to a conservative thresholding of density and/or streamlines to highlight the prevalence of this problem. Thus, these numbers are derived from a conservative estimate of the complexity of the brain, and represent the bare minimum of the number of convergent bundles. However, the absolute quantification itself may be biased towards the pathways from the chosen techniques and the atlases these are based upon. Additionally, while the bundles defined by these techniques have proven reliability, they are not themselves immune to the problems posed by bottlenecks. It is possible that improvements in tracking will refine the definitions of the bundles, as well as identify new pathways; while this may change estimates of the number of bundles in a given fixel, it won’t eliminate the basic feature of the coincidence of bundles in common pathway segments.

### Adding to known atlases

While the primary aim of this study was to characterize bottlenecks, in which >1 bundle passes through a fixel, we made the interesting ancillary observation (Supplementary Figure 1) that many fixels within the white matter were not associated with any bundles defined in our utilized atlases. We note that these ‘zero-bundle’ fixels were not included in the analysis because there is a lack of information regarding these orientations. While many of these could be spurious peaks or isolated voxels in white matter, larger expanses of coherently oriented FODs could be regions which may represent underexplored white matter pathways which can eventually be added to our repertoire and collection of bundles. While some regions are easily explained, for example due to a lack of TractSeg bundles in the cerebellum, other regions occur both along and across several gyral blades, as well as the relatively underexplored system of U-fibers and local association fibers. Further reasons for this could be the thresholding of densities and streamlines and parameter configurations for the bundle segmentation techniques. Additional atlases or bundle segmentation procedures may include pathways through many of these regions, which would solve the missing-bundle problem, but would likely also increase the prevalence of bottlenecks.

This is a potential limitation of the current study – the choice of atlases (bundle segmentation procedures). We have purposefully chosen to only include pathways for which there is broad agreement on their existence, and techniques which have been validated and well-utilized in the field. There would also be the potential to define pathways and bottleneck regions using whole-brain connectivity, or atlases which are derived from clusters or reproducibility of large datasets [48–50], however, these are potentially confounded by the bottleneck problem itself (i.e., contain false positive, yet reproducible, pathways), and an extensive comparison of algorithms and segmentation methods is beyond the scope of the current study.

### “Single Fiber Populations” and microstructure

In addition to tractography, the diffusion signal that results from a single fiber population (i.e., the fiber response function) has applications towards tissue microstructural modeling. The response function can be used to estimate the FOD and tissue microstructural properties including diffusivities, compartment sizes, and orientation dispersions. Typically, this response function is derived from studying regions of low complexity, specifically, regions that are considered single fiber populations and contain only a single peak in the FOD [25, 32, 51]. Here, however, we can see that even if a voxel or region contains a dominant orientation and high anisotropy, a majority of these regions are composed of multiple, distinct fiber pathways, that may have varying densities, sizes, and distributions of axons. For example, if we were to use the traditional definition of a ‘single fiber population’ (i.e., our single-orientation voxel) we would find that only 35% and 27% contain just a single fiber bundle passing through them (Supplementary Figure 2). Thus, these so-called single fiber regions are very often multi-bundle regions. Truly single-fiber and single-bundle regions are rare, even with our limited selection of known bundles used in this study, and in our case, occur in the cingulum and specific gyral blades (Supplementary Figure 3). While some works show that biological differences between fiber populations are negligible in the response function formulism [52], there is some evidence that the response functions do vary across pathways [18, 53, 54], which may lead to variation in estimates of FODs and subsequent microstructure.

### Nomenclature

Here, we have also introduced slightly different nomenclature than past literature. As described above a single-orientation voxel has traditionally been called a single-fiber voxel, whereas a multi-orientation voxel has been called crossing-fiber voxel. Clearly, a voxel with only one orientation is not limited to only containing the presence of fibers from a single bundle, hence the new description. Additionally, a fixel traditionally refers to a “specific fiber bundle within a specific voxel” [29, 55–57], yet we have shown again that a fixel (which is truly a segmented lobe, or orientation, from the FOD), is also not limited to a single specific fiber bundle, and in fact, likely contains axons from several bundles. We are not proposing to change the use and discussion of these elements throughout the field, but rather chose clarifying nomenclature to remove ambiguity in this specific study.

## Conclusion

In this work, we investigated the prevalence of bottleneck regions, or where multiple white matter pathways of the brain converge and subsequently diverge. Our results indicate that most white matter contains multiple overlapping bundles, and individual orientations within a voxel are associated with multiple bundles. These findings have profound implications for tractography analysis which aims to map unknown connections across the brain, and strengthen the awareness of limitations or challenges facing these image processing techniques.

## Data Availability

All resulting voxel-wise and fixel-wise bundle overlaps, for both TractSeg and Recobundles, is available upon request.

## Acknowledgments

This work was supported by the National Institutes of Health under award numbers R01EB017230, and T32EB001628, and in part by ViSE/VICTR VR3029 and the National Center for Research Resources, Grant UL1 RR024975-01. CMWT was supported by a Sir Henry Wellcome Fellowship (215944/Z/19/Z) and a Veni grant (17331) from the Dutch Research Council (NWO). MD was supported by the institutional Research Chair in Neuroinformatics and NSERC Discovery.

## Appendix

The bundles resulting from each bundle-segmentation pipeline are given as a list below.

### Recobundles

Arcuate Fasciculus left; Arcuate Fasciculus left; Frontal Aslant Tract left; Frontal Aslant Tract right; Cerebellum left; Cerebellum right; Corpus Callosum Major; Corpus Callosum Minor; Central Tegmental Tract left; Central Tegmental Tract right; Extreme Capsule left; Extreme Capsule right; Fronto-pontine tract left; Frontopontine tract right; Inferior Fronto-occipital Fasciculus left; Inferior Fronto-occipital Fasciculus right; Inferior Longitudinal Fasciculus left; Inferior Longitudinal Fasciculus right; Middle Cerebellar Peduncle; Middle Longitudinal Fasciculus left; Middle Longitudinal Fasciculus right; Medial Longitudinal fasciculus left; Medial Longitudinal fasciculus right; Medial Lemniscus left; Medial Lemniscus right; Occipito Pontine Tract left; Occipito Pontine Tract right; Optic Radiation left; Optic Radiation right; Parieto Pontine Tract left; Parieto Pontine Tract right; Superior longitudinal fasciculus left; Superior longitudinal fasciculus right; Spinothalamic Tract left; Spinothalamic Tract right; Temporopontine Tract left; Temporopontine Tract right; Uncinate Fasciculus left; Uncinate Fasciculus right; Vermis.

### TractSeg

Arcuate fascicle left; Arcuate fascicle right; Anterior Thalamic Radiation left; Thalamic Radiation right; Commissure Anterior; Rostrum; Genu; Rostral body (Premotor); Anterior midbody (Primary Motor); Posterior midbody (Primary Somatosensory); Isthmus; Splenium; Corpus Callosum – all; Cingulum left; Cingulum right; Corticospinal tract left; Corticospinal tract right; Frontopontine tract left; Fronto-pontine tract right; Fornix left; Fornix right; Inferior cerebellar peduncle left; Inferior cerebellar peduncle right; Inferior occipito-frontal fascicle left; Inferior occipito-frontal fascicle right; Inferior longitudinal fascicle left; Inferior longitudinal fascicle right; Middle cerebellar peduncle; Middle longitudinal fascicle left; Middle longitudinal fascicle right; Optic radiation left; Optic radiation right; Parieto-occipital pontine left; Parietooccipital pontine right; Superior cerebellar peduncle left; Superior cerebellar peduncle right; Superior longitudinal fascicle III left; Superior longitudinal fascicle III right; Superior longitudinal fascicle II left; Superior longitudinal fascicle II right; Superior longitudinal fascicle I left; Superior longitudinal fascicle I right;Striato-fronto-orbital left; Striato-fronto-orbital right; Striato-occipital left; Striato-occipital right; Striato-parietal left; Striato-parietal right; Striato-postcentral left; Striato-postcentral right; Striato-precentral left; Striato-precentral right; Striato-prefrontal left; Striato-prefrontal right; Striato-premotor left; Striato-premotor right; Superior Thalamic Radiation left; Superior Thalamic Radiation right; Thalamo-occipital left; Thalamo-occipital right; Thalamo-parietal left; Thalamo-parietal right; Thalamo-postcentral left; Thalamo-postcentral right; Thalamo-precentral left; Thalamo-precentral right; Thalamo-prefrontal left; Thalamo-prefrontal right; Thalamo-premotor left; Thalamo-premotor right; Uncinate fascicle left; Uncinate fascicle right.

