## Supplementary for "Prevalence of white matter pathways coming into a single diffusion MRI voxel orientation: the bottleneck issue in tractography"

**Supplementary Data**

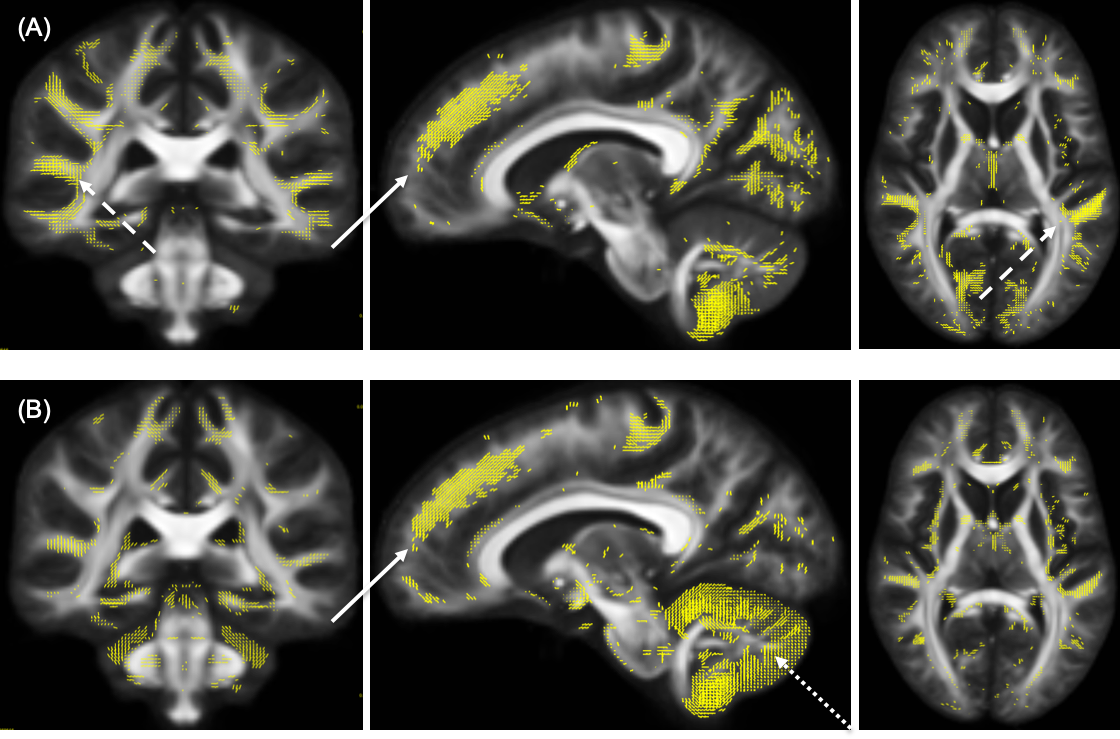

**Supplementary Figure 1.** Template space fixels in which zero fiber bundles are observed for Recobundles (A) and TractSeg (B).

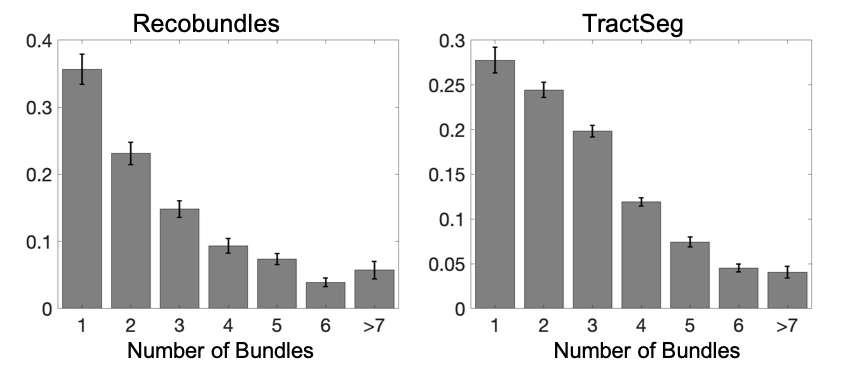

**Supplementary Figure 2**. Within voxels that only have one dominant orientation (i.e., one fixel), most have greater than one known bundle passing through that voxel. Bar plots show the number of bundles assigned to single fixel voxels for Recobundles and TractSeg algorithms.

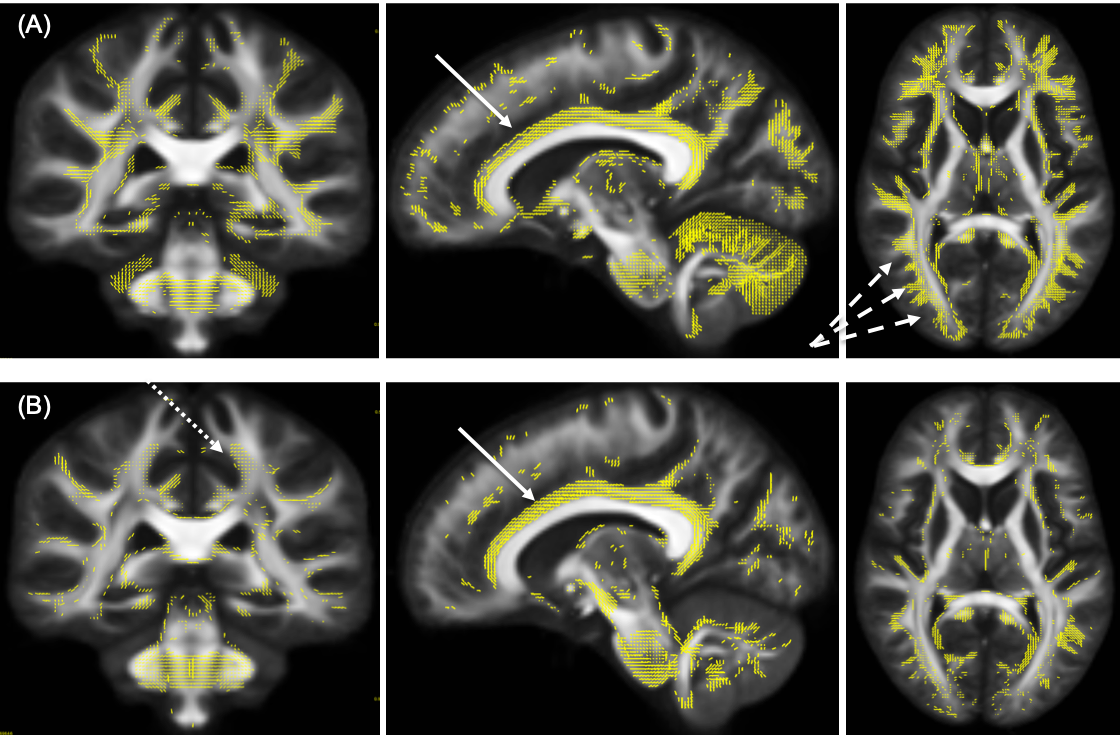

**Supplementary Figure 3.** Template space fixels in which a single fiber bundle is observed for Recobundles (A) and TractSeg (B).
